# Muscle-fiber specific genetic manipulation of Drosophila sallimus severely impacts neuromuscular development, morphology, and physiology

**DOI:** 10.1101/2024.04.15.589595

**Authors:** Andrew H. Michael, Tadros A. Hana, Veronika G. Mousa, Kiel G. Ormerod

**Affiliations:** Middle Tennessee State University, Murfreesboro, Tennessee, United States of America

**Keywords:** Drosophila, neuromuscular junction, sarcomere, muscle, sallimus, elastic protein

## Abstract

The ability of skeletal muscles to contract is derived from the unique genes and proteins expressed within muscles, most notably myofilaments and elastic proteins. Here we investigated the role of the sallimus (*sls*) gene, which encodes a structural homologue of titin, in regulating development, structure, and function of *Drosophila melanogaster*. Ubiquitous knockdown of *sls* using RNA interference (RNAi) in all muscle fibers resulted in embryonic lethality. A screen for muscle-specific drivers revealed a Gal4 line that expresses in a single larval body wall muscle in each abdominal hemisegment. Disrupting *sls* expression in single muscle fibers did not impact egg or larval viability nor gross larval morphology, but did significantly alter the morphology of individual muscle fibers. Analysis of individual sarcomeres revealed significant changes in ultrastructural organization, dramatically increasing sarcomere, I-band, A-band, and *sls*-band lengths. Surprisingly, muscle-cell specific disruption of *sls* also severely impacted neuromuscular junction (NMJ) formation. The extent of motor-neuron (MN) innervation along disrupted muscles was significantly reduced along with the number of glutamatergic boutons, in MN-Is and MN-Ib. Electrophysiological recordings revealed a 40% reduction in excitatory junctional potentials correlating with the extent of motor neuron loss. Analysis of active zone (AZ) composition revealed changes in presynaptic scaffolding protein (brp) abundance, but no changes in postsynaptic glutamate receptors. Ultrastructural changes in muscle and NMJ development at these single muscle fibers were sufficient to lead to observable changes in neuromuscular transduction and ultimately, locomotory behavior.

## Introduction

Skeletal muscle enables animals to produce movement, facilitating a robust set of behaviors and interactions with the environment. The force necessary for movement is created by linking two rigid skeletal elements and pulling them together. The repeating, functional units of skeletal muscle, the sarcomere, are similarly organized, bordered by rigid, structural Z-discs [1,2]. Sarcomeric striations, first noted in whale tissue by van Leeuwenhoek in 1712, are composed of two antiparallel filament systems, with thin actin-filaments sliding on thick myosin-filaments to shorten the sarcomere [3]. H. Huxley proposed the crossbridge theory where cyclic interactions occurring between myosin-based crossbridges with specialized attachment points on actin-filaments generated muscle contraction and force production [4]. While the crossbridge theory of the sarcomere captures many features of contracting muscle, it does not predict or account for many experimentally observed properties in skeletal muscle from; what maintains the structural integrity of the contracting sarcomere, to the restoration of sarcomere length following crossbridge cycling, or the incredibly long-range elasticity observed in sarcomeres [5]. Indeed, Huxley himself recognized the insufficiencies of the crossbridge conception and notes a ‘special feature’ must have evolved to facilitate these structural and functional properties of muscle [6].

The giant filamentous protein titin, spanning half the sarcomere from Z-disc to M-band is now largely accredited as that special feature [7]. Titin is ubiquitously expressed in all skeletal muscle, is the third most abundant muscle protein (after actin and myosin), and the largest known protein [8,9]. Titin’s completed genomic sequence revealed it to be a molecular spring; within its I-band region are compliant proximal Ig-domains which straighten at low forces, and the stiffer PEVK region extends when substantial force is applied [10]. Titin’s role in contributing to both active force production and passive tension are well established [8,9]. As research into titin has steadily grown, countless novel putative roles have surfaced including: a substrate for calcium binding, target for proteostasis and posttranslational modifications (oxidation or phosphorylation), mediating the Blaschko effect, mechanosensory mechanism, signaling hub, muscle stiffness regulation, and structural assembly/organization of sarcomeres [8–13]. A great deal remains to be understood regarding the various suggested roles of titin, however, *in vivo* models have proven difficult given that genetic alteration in titin expression leads to embryonic lethality in most species [14–17].

From insects to humans the structure and function of the sarcomere is well conserved [18]. Given that most components of the sarcomere, including titin, are highly conserved among animals, insect models have flourished as an essential aspect of muscle research [19]. *Drosophila* has emerged as an excellent model for muscle development and structure-function investigations given their structural and genetic conservation along with the large number of accessible genetic, molecular, and physiological tools [20]. The two *Drosophila* homologues of titin are Sallimus (*sls*) and Projectin (gene named bent, *bt*). A recent study demonstrated that *sls* encoded a protein which spans the I-band, from Z-disc to M-line, and it contains immunoglobulin, fibronectin, and PEVK domains along with other critical structural components, and targets of modifications and protein-interactions [17,21,22]. However, even within *Drosophila*, investigations of *sls* are scarce, as genetic manipulations led to embryonic lethality [17,21].

Here we initially verified that ubiquitously reducing the expression of *sls* in muscle cells resulted in embryonic lethality. A genetic Gal4 screen identified a unique line with expression in a single muscle fiber within each abdominal hemisegment of larvae. Reducing *sls* expression in a reduced number of muscles circumvented lethality and facilitated a comprehensive examination of the roles of *sls* in the development, structure, and function of muscle. Immunohistochemical analyses of muscle revealed a dramatic change in the size and morphology of muscles with reduced *sls* expression. The change in muscle structure also resulted in a profound change in the shape and length of motor neuron innervation along the surface of the muscle, along with the number and ultrastructural composition of synaptic boutons. These neuromuscular changes significantly reduced neuromuscular transduction revealed by electrophysiological recordings, and ultimately manifested in significantly reduced locomotory behavior.

## Results

The *Drosophila* neuromuscular system is ideal for investigations of genes and proteins in regulating muscle development, structure, and function (Fig. 1). Adult flies mate to produce fertilized eggs which hatch and undergo 3 larval instar stages. Afterwards, the larvae undergo metamorphosis as pupae, to then become adults. Third-instar larvae have a well-defined neuromuscular system, with 36 genetically programmed motor neurons forming stereotypical connections to the 30 muscles found in each abdominal hemisegment (Fig. 1 A-C) [23]. Each muscle is a viscerally located, striated multinucleated muscle fiber, attached directed to the cuticle through apodemes, which can be examined at the cellular or ultrastructural level by immunohistochemistry (Fig. 1 D-G) [24,25]. Typical muscle contractions are elicited via activation of glutamate receptors following the release of glutamate from presynaptic motor neuron terminals [26]. Motor neuron activity, which controls the release of glutamate, is dictated by neuronal input from descending interneurons onto the motor neuron soma, located in the ventral nerve cord (VNC). Locomotory activity is ultimately controlled by central pattern generators within the VNC [26].

**Figure 1:**
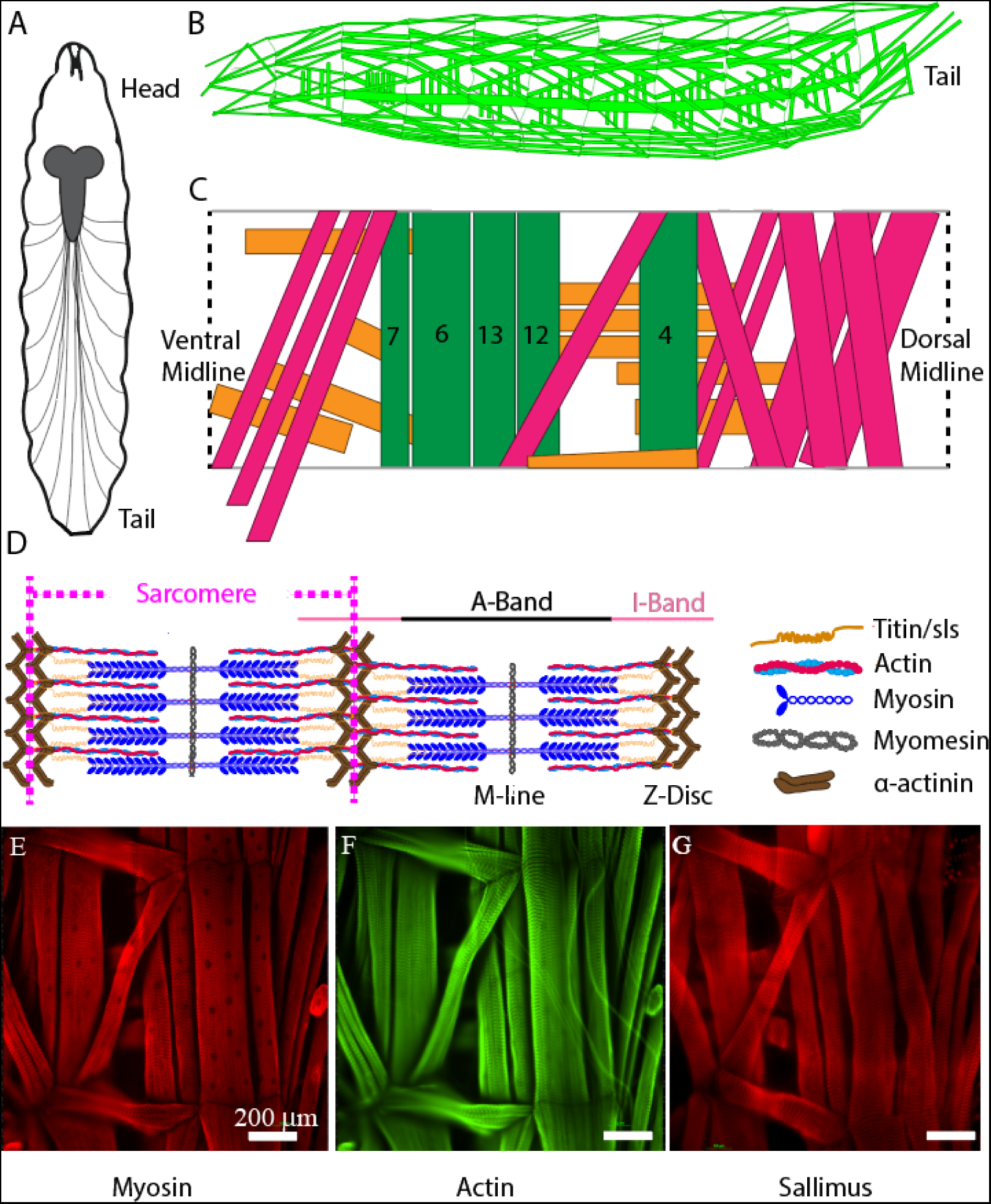
Schematic representation of critical sarcomeric proteins. ***A:*** Model of third-instar *Drosophila* larvae highlighting the CNS and main segmental nerve branches projecting to body-wall muscles. ***B:*** Three-dimensional representation of body-wall muscles in third-instar larvae. ***C:*** Schematic of a single abdominal right hemisegment highlighted in green, 5 commonly investigated longitudinal muscles which contribute most substantially to larval peristalsis. Orange and pink indicate transverse and oblique muscle respectively. ***D:*** Structure of a sarcomere highlighting 5 critical proteins involved in sarcomeric structure/function, as well as the A- and I-bands, M-line, and Z-disc. ***E-F:*** Immunohistochemical stains of third-instar body-wall muscles using anti-myosin ***E***, anti-actin ***F***, and anti-sallimus ***G***.

Recent work suggests that *Drosophila* sallimus (*sls)* encodes an elastic protein which spans from the sarcomeric Z-disc to myosin heads [17]. To examine the structural and physiological roles of *sls* in muscle, we took advantage of the genetic toolkit in *Drosophila* to alter its endogenous expression. A ubiquitous muscle driver (Mef2-Gal4, Fig. 2 A) was used to knockdown the expression of *sls* using RNAi. The number of eggs produced from 3-day old, mated females were tabulated from 2 different *sls*-RNAi lines (Fig. 2 B). No significant differences were observed compared to controls (Fig. 2 C, One-way ANOVA, F=2.03, P=0.62). The number of eggs that progressed to first-instar larvae were counted from each of the 6 genotypes in Fig. 2 C. From the Mef2-Gal4>UAS-sls1-RNAi 0 first-instars emerged from 150 eggs, and 3 first-instars emerged from 121 eggs from Mef2-Gal4>UAS-sls2-RNAi (Fig. 2 D, One-way ANOVA, F=29.69, P<0.0001). None of the larvae from our ubiquitous *sls* knockdowns progressed beyond first-instar, indicating embryonic lethality. Given that both UAS-sls-RNAi lines were equally effective, the remainder of our experiments were conducted with UAS-sls2-RNAi (hereto forth referred to as sls-RNAi). An additional ubiquitous muscle driver, MHC-Gal4, was crossed with our sls-RNAi line to validate the effects on larval lethality (Fig. 2 D). Collectively these data support previous findings that ubiquitous knockdown of *sls* in muscle fibers leads to early larval lethality [17,21].

**Figure 2:**
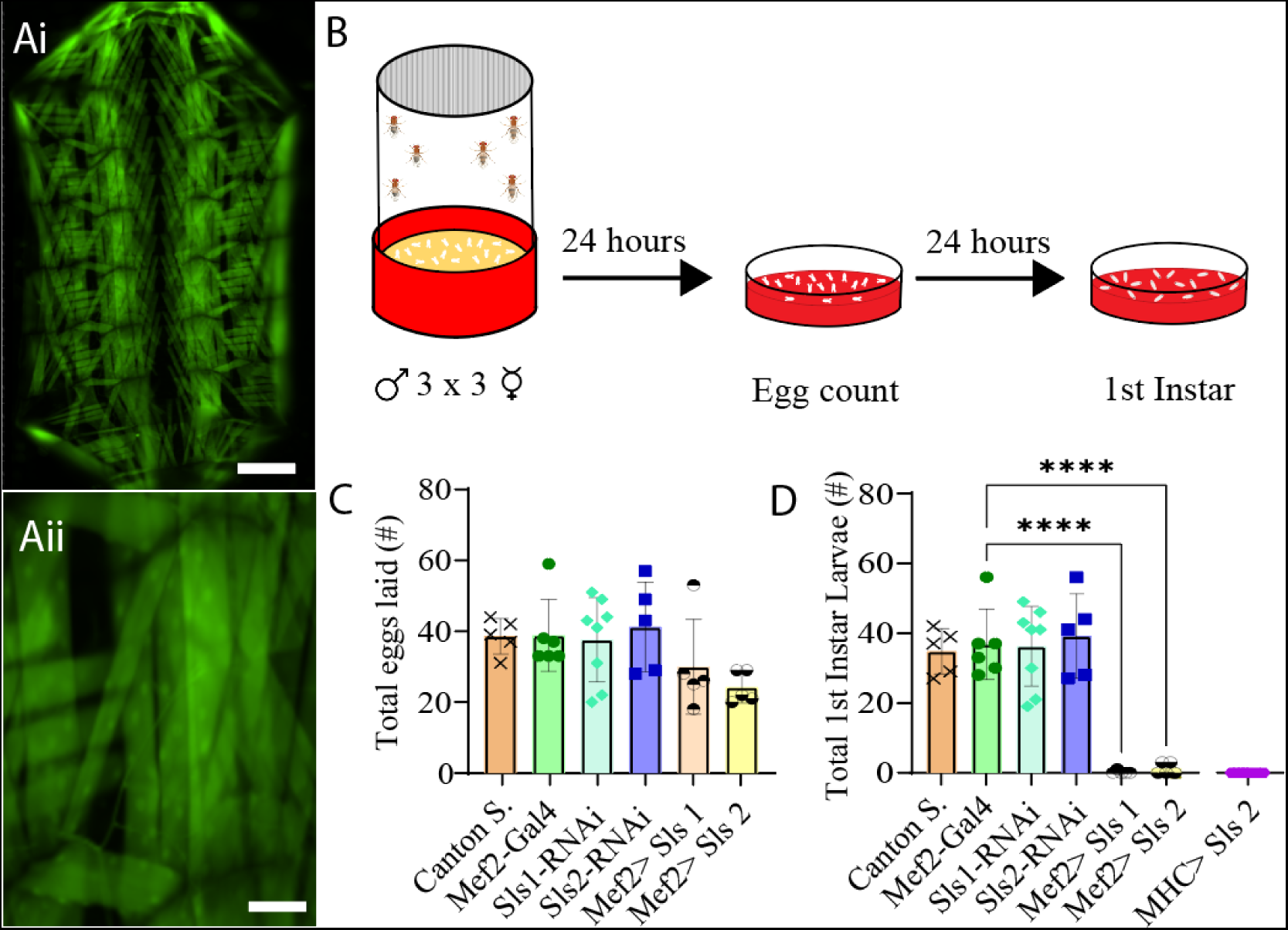
Ubiquitous RNAi knock-down of sallimus in body-wall muscles results in embryonic lethality. ***Ai:*** Fluorescence image of a filleted entire Mef2-Gal4>UAS-GFP third-instar larvae. Scale bar indicates 1000 μm. ***Aii:*** Fluorescence image of part of abdominal hemisegment A3 focused on MF 12. Scale bar indicates 175 μm. ***B:*** Schematic representation of the egg laying assay followed by egg and first-instar larvae counting after 24 and 48 hours respectively. ***C:*** Total eggs laid from the 6 genotypes investigated (N=5-8, One-way ANOVA, F= 2.03) ***D:*** Total first-instar larvae that emerged from the egg laid in ***C***, plus one additional muscle driver (myosin heavy chain, MHC) expressing Sls-RNAi (N=5-8, One-way ANOVA, F=29.69).

To circumvent lethality a genetic screen was conducted for Gal4-drivers that express in a subset of third-instar larval muscle fibers. Ten different muscle drivers were obtained from BDRC, crossed with UAS-Green Fluorescent Protein (GFP), and imaged using fluorescence microscopy. Of those lines, 3 exhibited GFP expression in a subset of muscle fibers; but 2 of which also had visible expression in the nervous system (NS). One line revealed expression *only* in muscle fiber 12, with no GFP expression in the NS (Fig. 3 B). This driver line was used in the remainder of this study and is subsequently referred to as MF12-Gal4. To determine if altering the expression of *sls* in only a subset of muscle fibers was sufficient to circumvent lethality, MF12-Gal4 was crossed with sls-RNAi. An egg-laying assay revealed no significant differences in total eggs laid between MF12-Gal4>sls-RNAi (MF12>sls) and controls (Fig. 3 D, One-way ANOVA, F=1.39, P=0.23). There was also no significant difference between the MF12>sls lines and the number of first-, second-, and third-instar larvae that emerged (only first-instar shown, Fig. 3 E One-way ANOVA, F=15.18, P>0.05). Next, 20 third-instar larvae were transferred to fresh food vials, and the percentage of larvae that enclosed to become pupae and subsequently adults were scored. Here again, no significant difference was observed in the pupal viability (Fig. 3 F, One-Way ANOVA, F=520.1, P>0.05) nor the number of adults that emerged from the initial 20 larvae (Fig. 3 G, One-way ANOVA, F=357.8, P>0.05).

**Figure 3:**
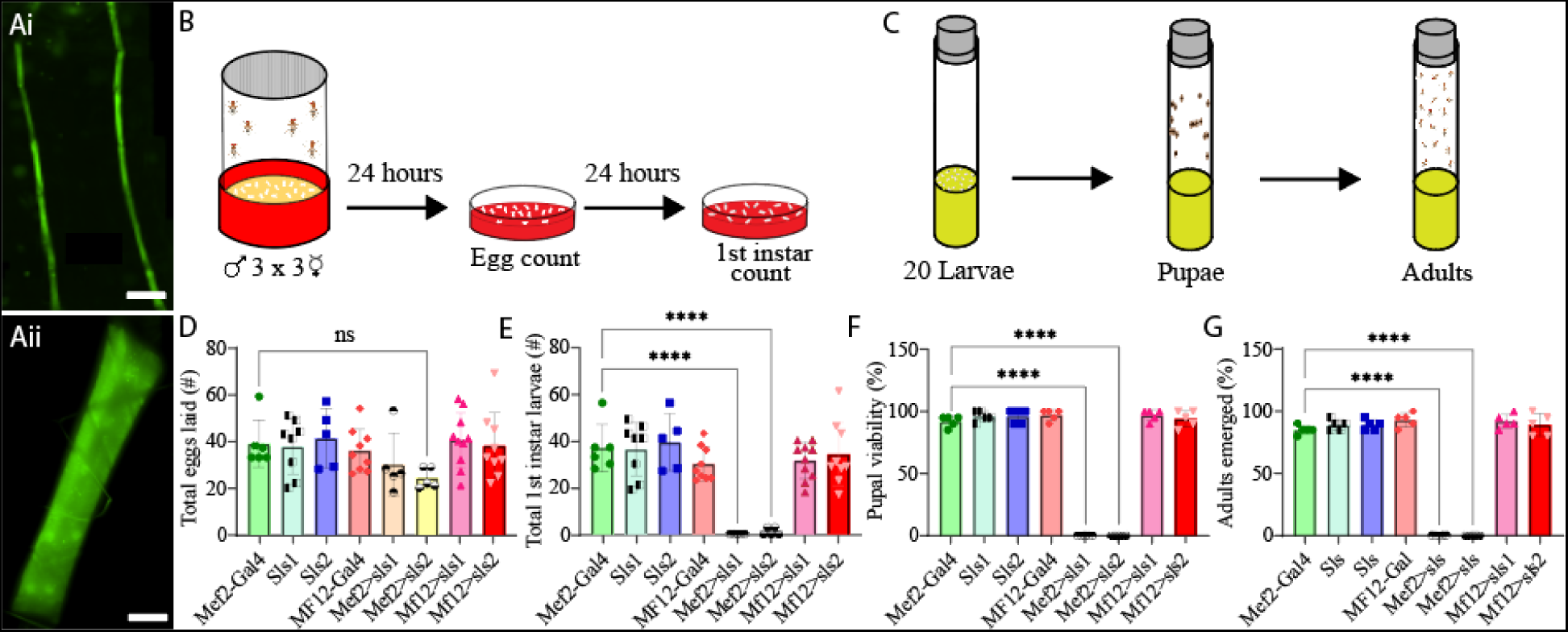
Muscle-fiber specific driver leads to larval, pupal, and adult viability. ***Ai:*** Fluorescence image of an immunohistochemical stain of an entire MF12-Gal4>UAS-GFP third-instar larvae. Scale bar indicates 800 μm. ***Aii:*** Immunostain image of part of abdominal hemisegment A3 focused on MFs 6, 7, 12, 13, and 4. Scale bar indicates 100 μm. ***B:*** Schematic representation of the egg laying assay followed by egg and first-instar larvae counting after 24 and 48 hours respectively. ***C:*** Schematic of pupal and adult viability assay. ***D:*** Total eggs laid from the 8 genotypes investigated (N=5-10, One-way ANOVA, F= 1.39, 0.22). ***E:*** Total first-instar larvae that emerged from the egg laid in ***D***, from the 8 genotypes investigated (N=5-10, One-way ANOVA, F=15.18, P>0.0001). ***F:*** Percent of 20 third-instar larvae that pupated (N=5, F=520.1, P<0.0001). ***G:*** Percent of 20 third-instar larvae that emerged to adulthood (N=5, F=357.8, P<0.0001).

Next, morphological metrics were obtained from third-instar larvae. No significant differences were observed for larval length, width, or area (Fig. 4, One-way ANOVA, F=2.13, P=0.95). To examine the role of *sls* in muscle structure, first, immunohistochemical analyses were conducted for actin (anti-phalloidin, Fig. 5 A-C). A dramatic gross structural change was observed in the morphology of MF 12 compared to controls (Fig. 5 A, C). The gross changes in muscle structure were first quantified by measuring the length, width, and area of MF 12 (Fig. 5 B). Significant differences were observed for muscle length (Fig. 5 Di, One-way ANOVA, F=20.2, P=0.0001), muscle width (Fig. 5 Dii, One-way ANOVA, F=368.0, P=0.0001), and muscle area (Fig. 5iii, One-way ANOVA, F=320.1, P=0.0001). The gross morphology of muscles 13 and 4 were also used as internal controls for all genotypes, and no significant differences were observed (Fig. 5 Ei-iii, Fi-iii).

**Figure 4:**
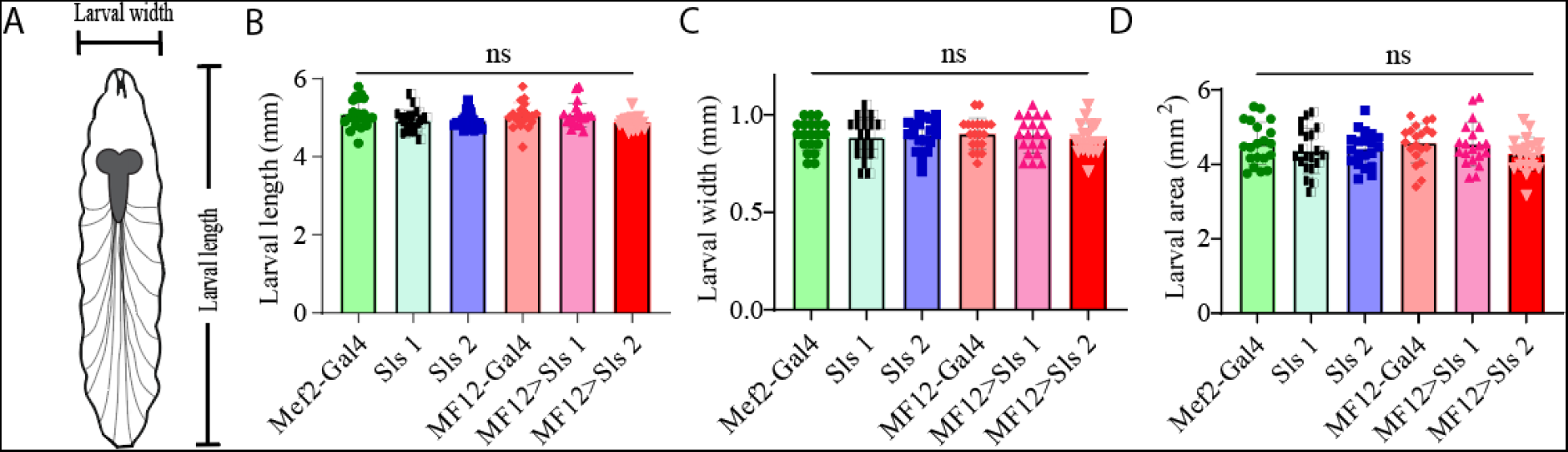
Muscle-specific sallimus knockdown does not alter larval morphology. ***A:*** Schematic representation of third-instar larvae indicating length and width measurements. ***B-D:*** Knocking down *sls* in MF12 did not significantly alter larval length (**B:** One-way ANOVA, F=2.13, P=0.66), width (**C:** One-way ANOVA, F=0.21, P=0.96), or area (**D:** One-way ANOVA, F=1.08, P=0.37).

**Figure 5:**
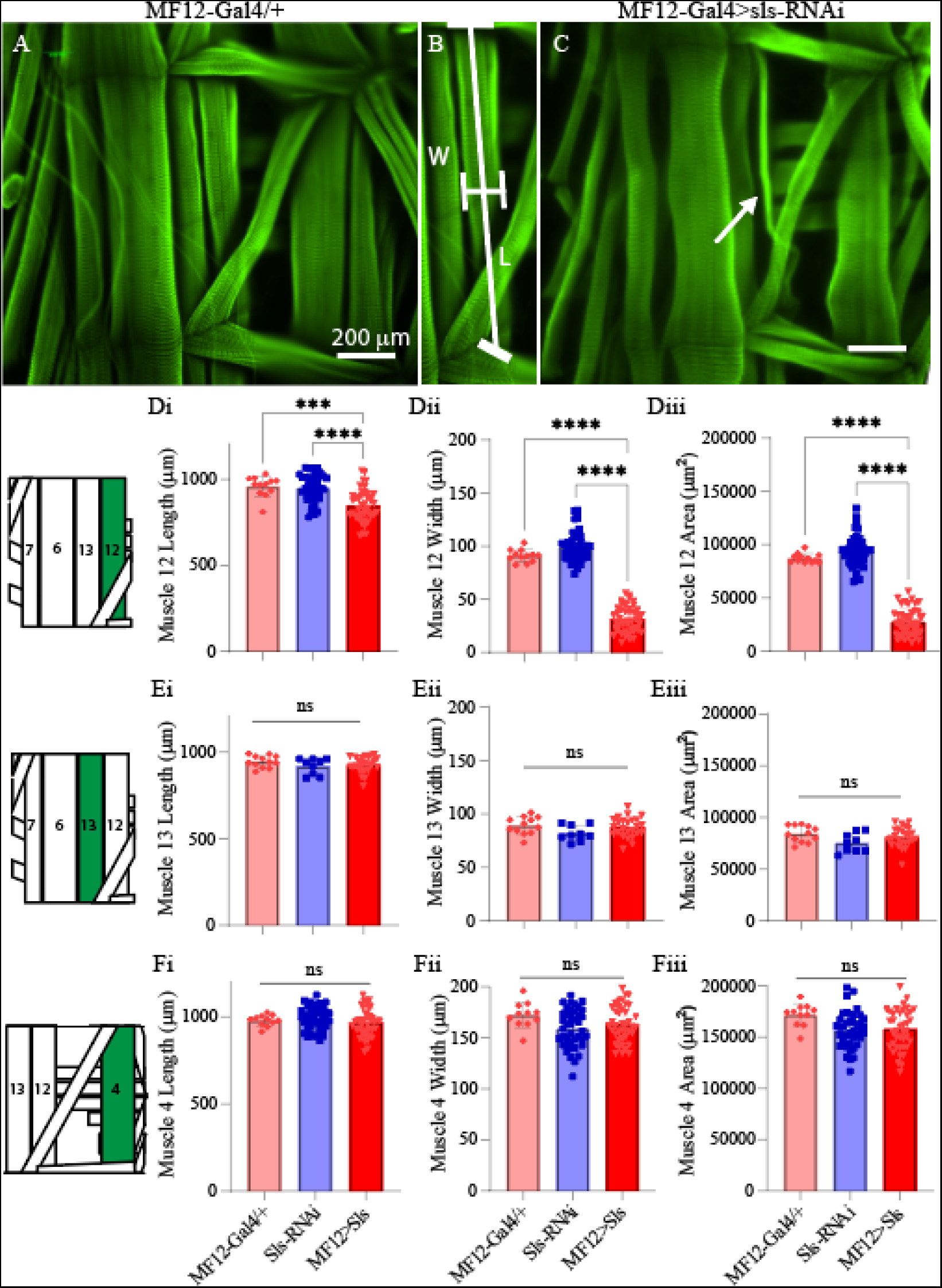
Muscle sallimus significantly impacts gross muscle morphology. ***A.*** Immunohistochemical stain for actin (phalloidin) in MF12-Gal4/+ larvae showing a single abdominal hemisegment. ***B.*** Immunohistochemical stain for actin highlighting muscle 12, along with indications for how length and width measurements were conducted. ***C.*** Immunostain from MF12-Gal4 > UAS-Sls-RNAi, white arrow indicates profound gross morphological change in MF12. ***D***: Quantification of muscle fiber 12 length (***Di***, One-way ANOVA, F=20.22, P<0.0001), width (***Dii***, One-way ANOVA, F=368.0, P<0.0001) and area (***Diii***, One-way ANOVA, F=320.1, P<0.0001). E: Quantification of muscle fiber 13 length (***Ei***), width (***Eii***) and area (***Eiii***). ***F***: Quantification of muscle fiber 4 length (***Fi***), width (***Fii***) and area (***Fiii***). Inset: cartoon depiction of abdominal larval segment, green highlights muscle being quantified.

Next, we established a novel approach to determine changes in MF ultrastructure by examining sarcomere, A-band, and I-band lengths (Fig. 6). Using a fluorescence line profile feature from Nikon Elements, calculations of both sarcomere and I-band length were made from animals immunostained with phalloidin (Fig. 6 Ai-ii). Individual sarcomere measurements were determined by measuring the distance between two phalloidin peaks (Fig. 6 Aii). Individual sarcomere lengths were 14.1 ± 0.2 µm and 14.4 ± 0.2 µm, for the control genotypes MF12-Gal4 and sls-RNAi, respectively (Fig. 6. Ei). Noticeably, MF12>sls animals demonstrated large variability in sarcomere lengths ranging from 19-to-47 µm, with an average of 31.7 ± 9.2 µm. I-band measurements also revealed a broad range from 16-33 µm with an average of 21.8 ± 5.5 µm, compared to 7.7 ± 0.1 µm and 7.8 ± 0.2 µm, for controls (Fig. 6 Eii: One-way ANOVA, F=100.9, P<0.0001). To examine changes in A-band measurements we co-stained animals with anti-myosin. A-band measurements showed an even greater range, from 16-27 µm with an average of 21.5 ± 4.3 µm, compared to controls 9.1 ± 0.2 µm and 9.2 ± 0.3 µm (Fig. 6 Eiii: One-way ANOVA, F=163.2, P<0.0001). Internal controls for sarcomere, I-band, and A-band measurements from muscles 13 and 4 were not significantly different from MF12 controls, nor between genotypes (Fig. 6 F-G). Collectively these results show a dramatic increase in sarcomere length and variability, and demonstrate incredible instability in the maintenance and formation of sarcomere assembly when *sls* expression is disrupted.

**Figure 6:**
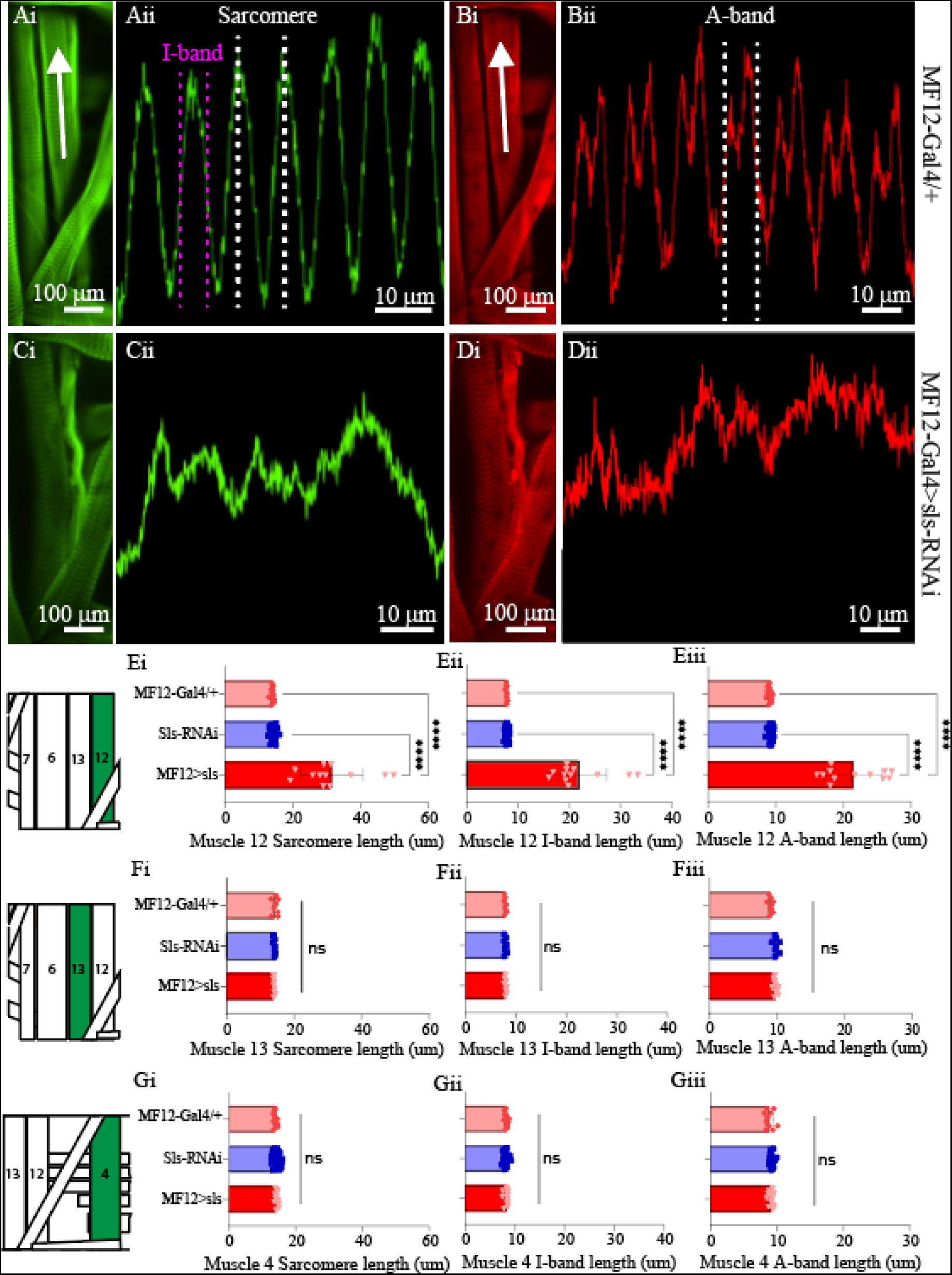
Muscle sallimus significantly impacts sarcomere assembly. Immunohistochemical stain for actin in **(*Ai*)** MF12-Gal4/+ and **(*Ci*)** MF12>Sls highlighting MF12. Arrow indicates location and direction of fluorescence intensity profile line for **(*Aii*)** and **(*Cii*)**. ***Aii:*** Fluorescence intensity profile line for GFP (actin). Dashed white and magenta lines indicate how sarcomere and I-band measurements were calculated respectively. Immunohistochemical stain for myosin in **(*Bi*)** MF12-Gal4/+ highlighting MF12 **(D*i*)** MF12>Sls highlighting MF12. Arrow indicates location and direction of fluorescence intensity profile line. ***Bii:*** Fluorescence intensity profile line for RFP (myosin). Dashed-line indicates a single myosin cycle, used for calculating A-band measurements. ***Dii:*** fluorescence intensity profile line for myosin from MF12>sls. ***E***: Quantification of muscle fiber 12 sarcomere length (***Ei***, One-way ANOVA, F=20.22, P<0.0001), I-band length (***Eii***, One-way ANOVA, F=368.0, P<0.0001) and A-band length (***Eiii***, One-way ANOVA, F=320.1, P<0.0001). ***F***: Quantification of muscle fiber 13 sarcomere length (***Fi***, One-way ANOVA, F=20.22, P<0.0001), I-band length (***Fii***, One-way ANOVA, F=368.0, P<0.0001) and A-band length (***Fiii***, One-way ANOVA, F=320.1, P<0.0001). ***G***: Quantification of muscle fiber 4 sarcomere length (***Gi***, One-way ANOVA, F=20.22, P<0.0001), I-band length (***Gii***, One-way ANOVA, F=368.0, P<0.0001) and A-band length (***Giii***, One-way ANOVA, F=320.1, P<0.0001).

To directly examine changes in *sls* protein ultrastructure, an immunohistochemical analysis for *sls* was conducted, which revealed rhythmic striations, emblematic of striated MFs (Fig. 7 A, B). Vibrant profile lines were observed for *sls* stained muscle fibers from control animals, and fluorescence intensity plots generated rhythmic peaks with an average width value of 9.1 ± 0.3 µm for MF12-Gal4, and 9.2 ± 0.3 µm for sls-RNAi. Sls immunostains of MF12>sls showed reduced fluorescence, and like the I- and A-bands, incredible variability in rhythmic striations ranging from 16-to-27 µm with an average of 21.5 ± 4.3 µm (Fig. 7 C-D). To quantify the reduction in *sls* protein expression, the average fluorescence intensity of MF12 area was calculated for each genotype, along with MF 13 to serve as an internal control. Average fluorescence for MF12 was 2641.3 ± 695.1 AU for MF12-Gal4, and 3074 ± 395.4 AU for sls-RNAi. For MF12>sls, average fluorescence of MF12 was 1058.9 ± 455.5, resulting in a significant reduction of 59 and 65% compared to the two controls (Fig. 7 E, One-way ANOVA, F=31.0, P<0.0001). This data confirms our sls-RNAi significantly reduces *sls* levels at MF12. Fluorescence intensity for MF13 was not significantly different between the three genotypes (data not shown).

**Figure 7:**
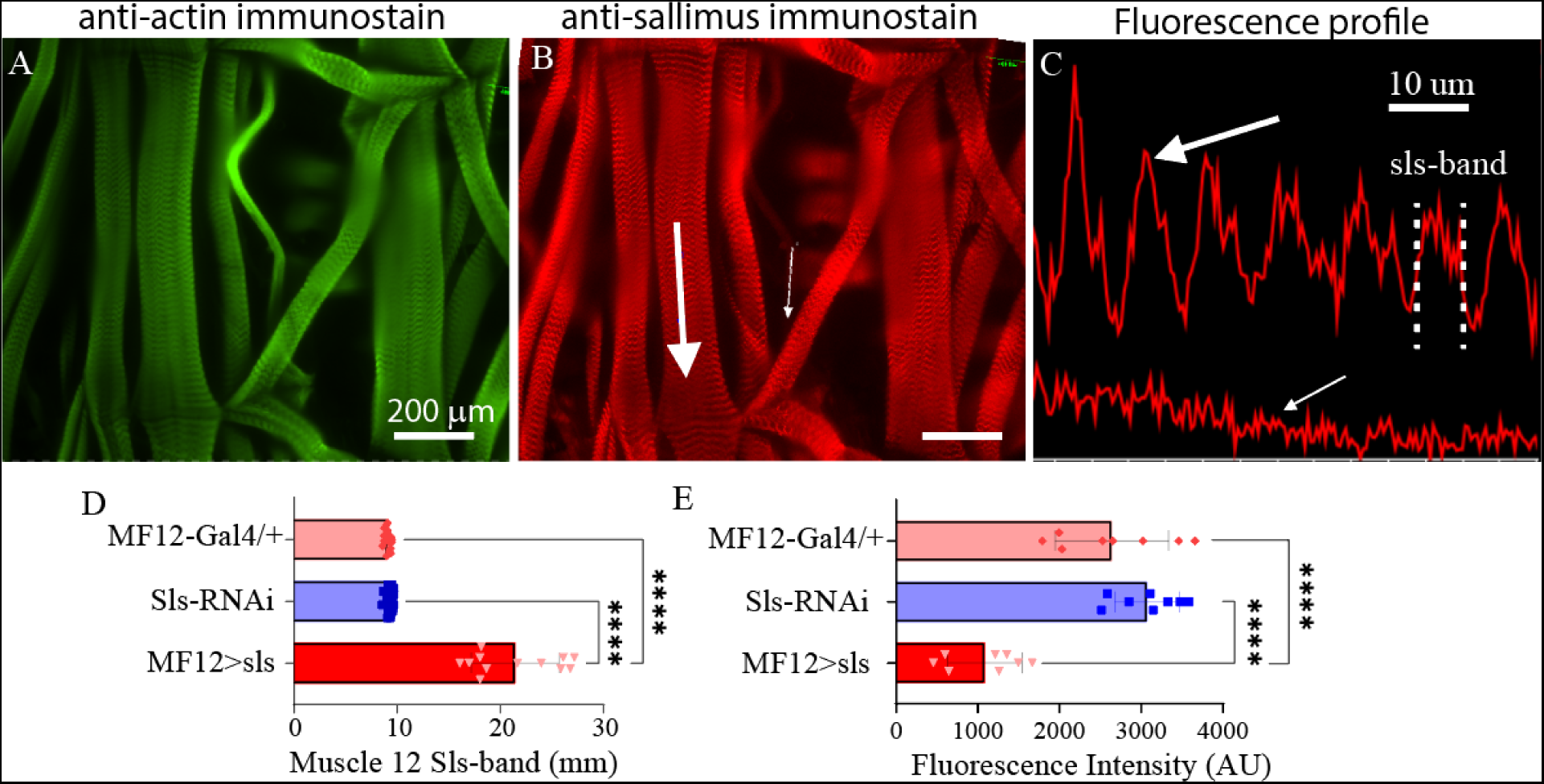
Reduced sls expression significantly increases length of sls-band. Phalloidin stain **(*A*)** and anti-sls from MF12>sls **(*B*)**. Large white arrow in **(*B*)** reveals location for fluorescence profile line in **(*C, top*)** from muscle 6, and small white arrow for muscle 12 **(*C*, *bottom*). (*D*)** Quantification of length of sls-band from 2 control and transgenic genotypes. **(*E*)** Sallimus fluorescence quantification from area of MF12 from all 3 genotypes.

A previous study using *Drosophila* noted that embryonic lethal mutations in *sls* resulted in malformations of multi-nucleate syncytia in embryonic muscles [21]. The nuclear stain DAPI was used to examine changes in nuclei number and intensity (Fig. 8). The average number of nuclei was 10.8 ± 0.4 for MF12-Gal4, 11.7 ± 1.0 for UAS-sls-RNAi, and 3.9 ± 0.9 for MF12>sls, demonstrating a 64 and 67% reduction in nuclei number in the *sls* knockdown flies compared to controls (Fig. 8 Ci, One way ANOVA, F=360.3, P<0.0001). The mean area of DAPI was also significantly reduced in MF12>sls MF 12, showing a 61% reduction compared to controls (Fig. 8 Cii, 488.5 ± 24, 433 ± 17, 188 ± 9, MF12-Gal4, sls-RNAi, MF12>sls respectively; one-way ANOVA, F= 101.2, P<0.0001). The mean DAPI fluorescence was significantly increased in MF12>sls muscle 12, on average showing 3.5 times greater fluorescence intensity compared to controls (Fig. 8 Ciii, 1400 ± 154, 3410 ± 198, and 8514 ± 577, MF12-Gal4, sls-RNAi, MF12>sls respectively, one-way ANOVA, F= 65.33, P<0.0001).

**Figure 8:**
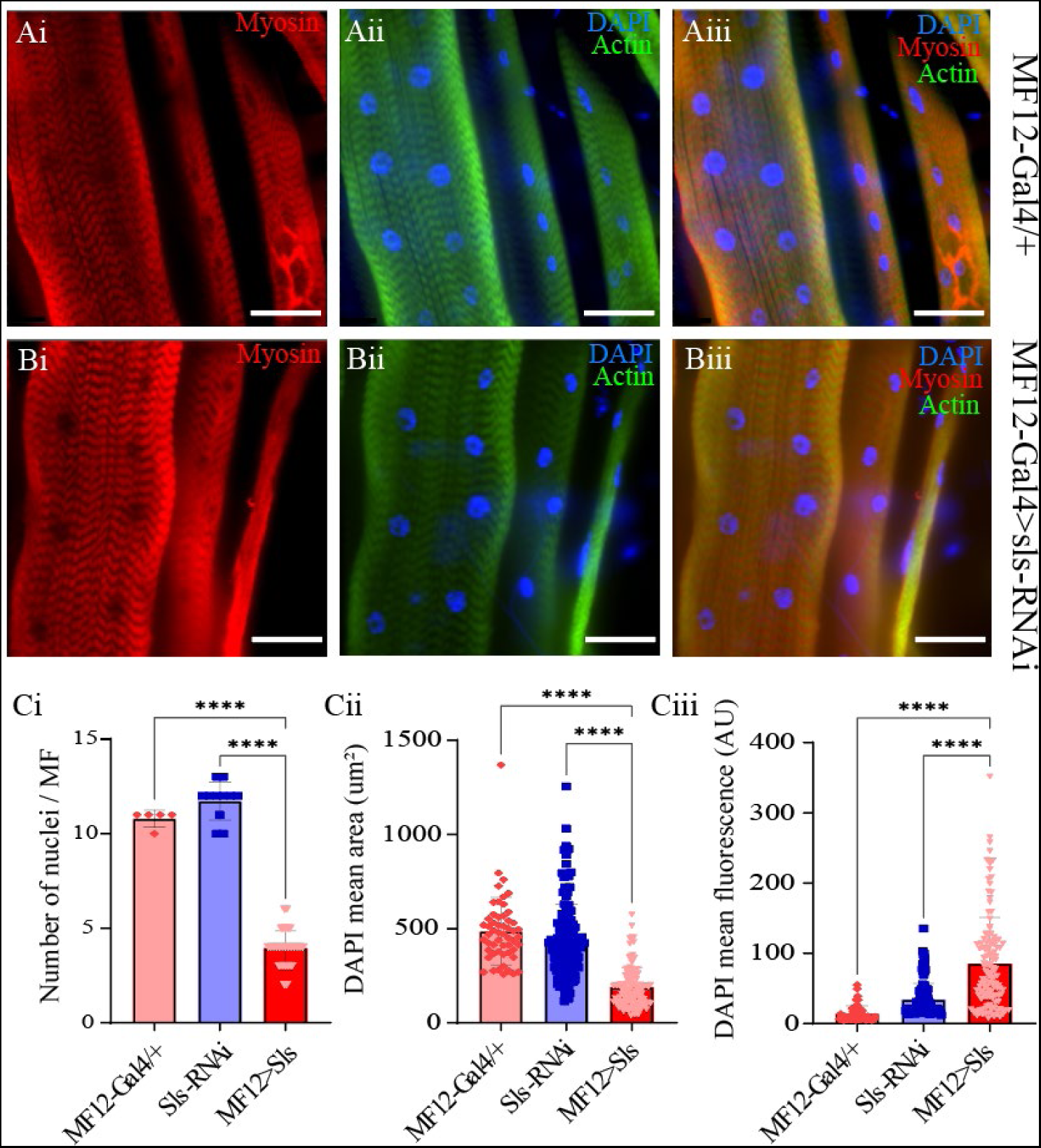
Sallimus disruption significantly reduces the number and size of muscle nuclei. Representative immunostains from MF12-Gal4/+ **(*Ai-Aiii*)** and MF12>sls **(*Bi-Biii*),** scale bars 100 µm. Blue: DAPI, green: anti-phalloidin red: anti-myosin. **(*Ci*)** Quantification of the number of nuclei per MF12. **(*Cii*)** Quantification of DAPI area from all DAPI-positive puncta from MF12. **(*Ciii*)** Quantification of DAPI fluorescence intensity from puncta in **(*Cii*)**.

A striking observation was made during the muscle ultrastructural analysis; the composition of MF12 NMJ was severely impacted when *sls* expression was reduced (Fig. 9). The extent of innervation as well as the size and morphology of glutamatergic MN terminals appeared to be severely affected (Fig. 9 A). *Drosophila* larvae have two different types of glutamatergic MN terminals, MN-Ib and MN-Is, comparable to mammalian tonic and phasic terminals, respectively. We first quantified the impact of *sls* disruption on NMJ morphology by examining the extent of innervation along the surface of each of these MN subtypes separately, then summated them together to generate a total (Fig. 9, Total, Ib, and Is rows). The length of NMJ innervation of MN-Ib was reduced by over 50% in MF12>sls compared to the two controls (Fig. 9 Cii, MF12-Gal4: 269.0 ± 67.0, sls-RNAi: 241.9 ± 68.8, MF12>sls: 124.2 ± 27.1, One-way ANOVA, F=14.73, P<0.0001). The length of innervation MN-Is were reduced by nearly 60% in MF12>sls compared to controls (Fig. 9 Ciii, MF12-Gal4: 313.0 ± 86.1, sls-RNAi: 319.4 ± 78.5, MF12>sls: 135.0 ± 38.4, One-way ANOVA, F=19.02, P<0.0001). Combining the two showed a 55% reduction in total innervation length (Fig. 9 Ci). Next, we assessed how disrupting *sls* impacted the total number of boutons per MN subtype innervating MF12. A 55% reduction in MN-Ib boutons was observed compared to controls (Fig. 9 Dii, MF12-Gal4: 15.0 ± 2.5, sls-RNAi : 15.4 ± 4.3, MF12>sls: 6.8 ± 1.3, One-way ANOVA, F=13.34, P<0.0009), and a 61% reduction in MN-Is boutons (Fig. 9 Diii, MF12-Gal4: 29.6 ± 7.3, sls-RNAi: 28.6 ± 13.1, MF12>sls: 11.2 ± 3.4, One-way ANOVA, F=13.34, P<0.0108). The combined total of 58% reduction in bouton size was observed for MF12>sls compared to controls (Fig. 9 Di). To examine changes in bouton ultrastructure more thoroughly, an assessment of active zone (AZ) composition was initially conducted using an immunostain against bruchpilot (brp), a ubiquitous presynaptic active zone (AZ) scaffold protein (Fig. 9 B). No changes in brp-positive puncta were observed (Fig. 9 Ei). However, significantly lower brp fluorescence intensity per puncta was observed for MN-Is and MN-Ib boutons in MF12>sls lines compared to controls (Fig. 9 Eii-Eiii, One-way ANOVA: MN-Is: F=22.67, P<0.0001; MN-Ib: F= 27.53, P<0.0001). Next, we examined postsynaptic ultrastructure via an immunostain for glutamate receptor III (GluRIII), one of the core subunits of postsynaptic glutamate receptors (Fig. 9 A). Neither GluRIII density nor fluorescence intensity was significantly different between MF12-Gal4 and MF12>sls for either MN-terminal subtype (Fig. 9 G-H).

**Figure 9.**
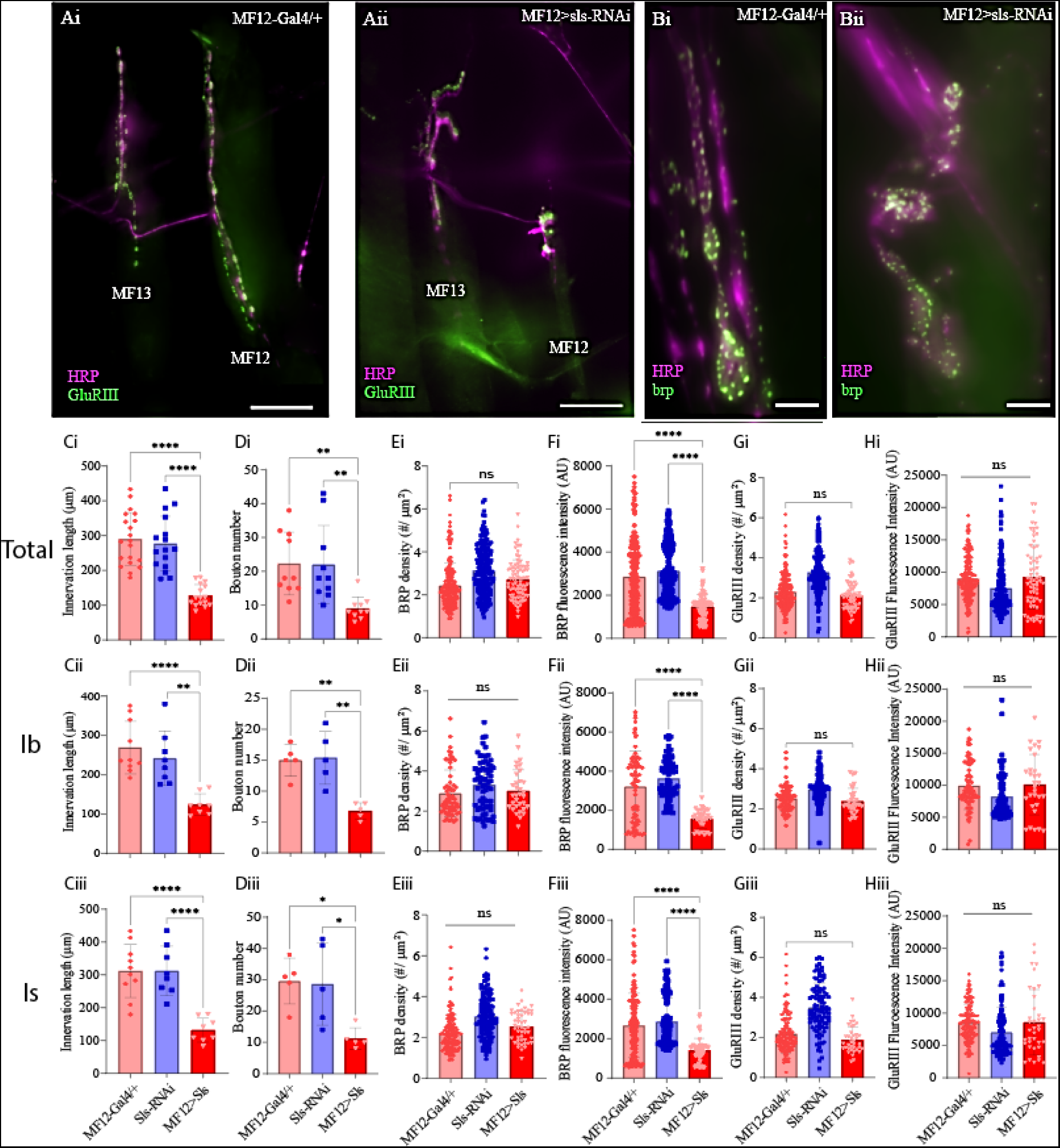
Disruption of muscle sls significantly impairs NMJ formation. ***A***: Immunohistochemical image of MN innervation along MF12 of a third-instar abdominal hemisegment with emphasis on muscles 12 and 13 from MF12-Gal4/+ (***Ai***) and MF12>sls (***Aii***), scale bar, 100 µm. Immunostain images for the AZ-scaffold protein brp highlighting MF12 from MF12-Gal4/+ (***Bi***) and MF12>sls (***Bii***), scale bar, 15 µm. Quantification of the impacts of *sls* disruption on synaptic innervation and NMJ formation (***Ci-Hiii***). Data are in rows presented based on total innervation (i), or broken down by MN subtype, MN-Ib (ii), MN-Is (iii). Changes in total innervation (***Ci-Ciii***) and bouton number (***Di-Diii***) from an HRP stain. Results from brp immunostaining depict AZ density (***Ei-Eiii***, brp density), and brp fluorescence intensity (***Fi-Fiii***). Examining postsynaptic NMJ changes using a GluRIII stain for changes in GluR density (***Gi-Giii***) and fluorescence intensity (***Hi-Hiii***).

Given the pronounced changes in MN and muscle ultrastructural, sharp intracellular electrophysiological recordings were taken from MF 12 and MF 6. Excitatory junctional potentials (EJPs) were elicited at low frequency stimulation (0.2 Hz) for 5 minutes. A significant, 40% reduction in the amplitude of EJPs was observed for MF12>sls compared to controls (Fig. 10, MF12-Gal4: 28.25 ± 3.1, sls-RNAi: 28.96 ± 4.8, MF12>sls: 16.01 ± 2.9, Canton S.: 26.7 ± 2.6, One-Way ANOVA, F=16.19, P<0.0001). As an internal control, EJPs were also recorded from muscle fiber 6, and no significant differences were observed (Fig. 10).

**Figure 10:**
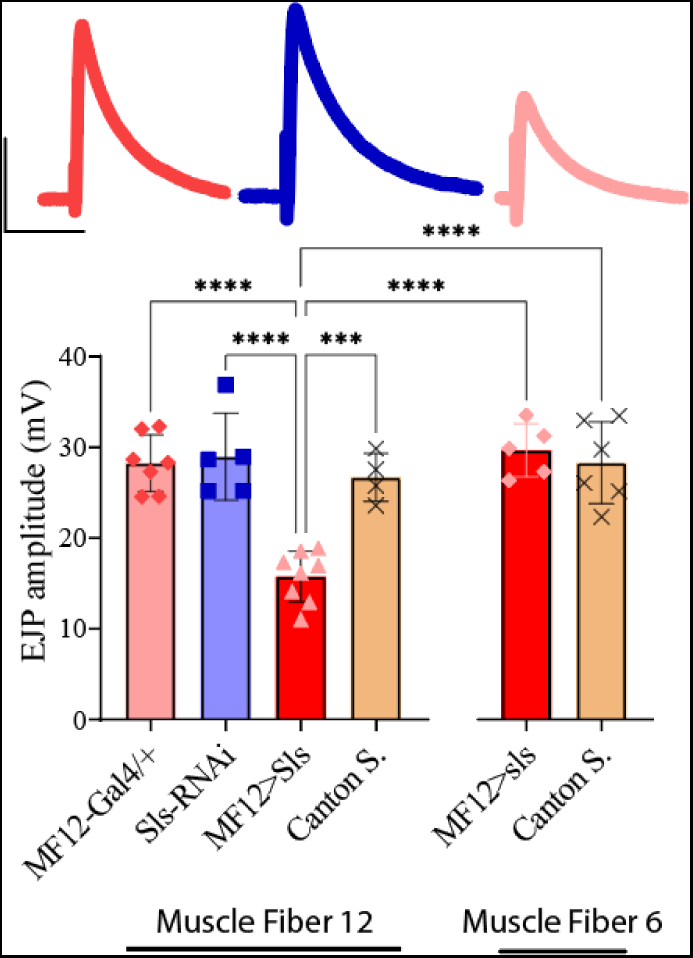
Excitatory junctional potentials are significantly reduced in sls knockdown muscles. EJP amplitudes averaged across each genotype show a significant reduction in MF12>sls compared to controls. Insets above from left to right: representative EJP traces from MF12-Gal4/+, UAS-sls-RNAi, and MF12>sls. Scale bars: 15 mV, 40 ms.

Neuromuscular transduction is an integral component of the circuitry underlying rhythmic peristalsis for larval locomotion. Given the changes in NMJ and muscle ultrastructure, we next assessed whether a behavioral change was observable following *sls* disruption. A larval crawling assay was conducted to assess changes in velocity, distance travelled, displacement, and angular velocity of third-instar larvae. One hundred larvae from each genotype were tracked for 5 minutes in a dark box using an infrared camera. The average velocity for controls was 0.88 ± 0.2 µm/ s for MF12-Gal4/+, 0.80 ± 0.2 µm/ s for sls-RNAi. The velocity for MF12>sls was reduced significantly by 25% compared to controls (Fig. 11, MF12>sls: 0.7 ± 0.2 µm/ s, One way ANOVA, F=11.2, P<0.0001). The distance traveled was also significantly reduced in MF12>sls larvae compared to controls (data not shown, One-way ANOVA, F=4.3, P<0.01). Angular velocity provides a metric to assess how much a larva turns during the recording period [27]. This was not significantly different between the three genotypes (data not shown). Taken together, these data reveal that, despite MF12-Gal4 expressing in a single muscle fiber within each hemisegment, disruptions in *sls* are sufficient to significantly reduce locomotory, larval crawling behavior.

**Figure 11.**
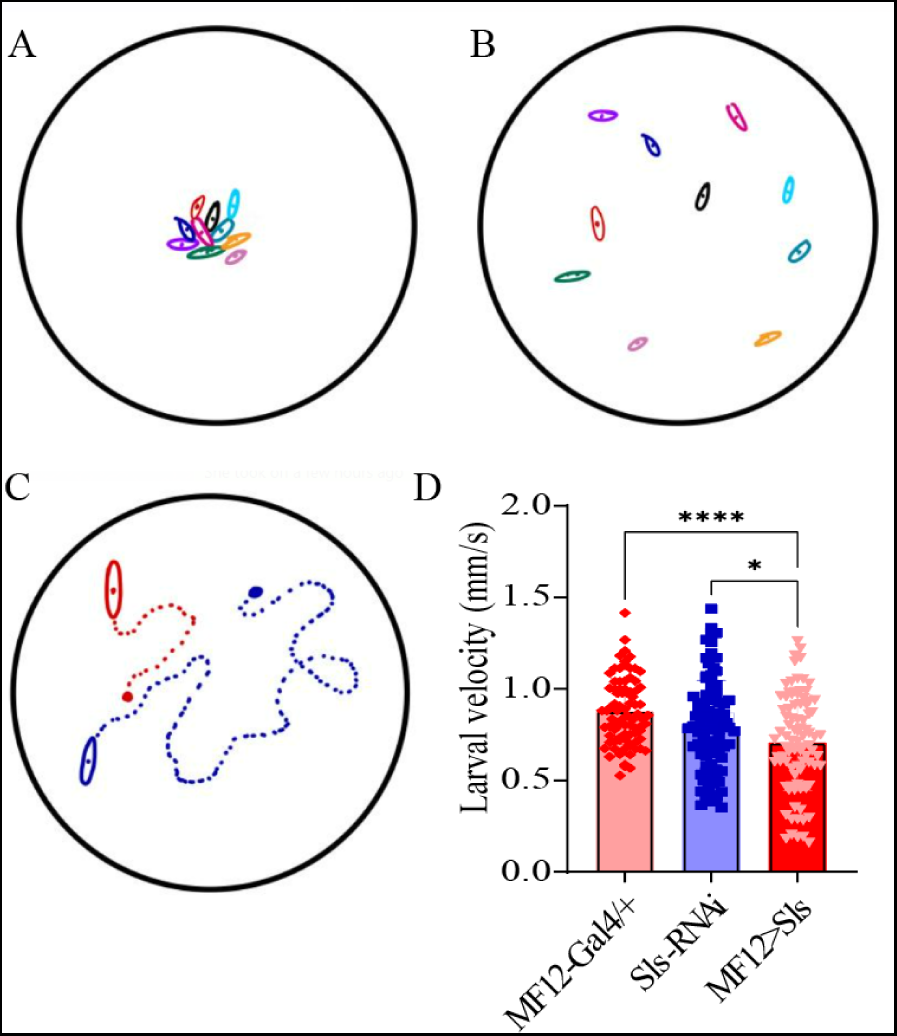
Muscle-fiber specific reduction in sls expression is sufficient to impair larval crawling behavior. ***A-C***: Schematic representations of 10 third-instar larvae placed in the center of an agar-bottomed-Petri dish (***A***), allowed to crawl for 5 minutes (***B***), and their position is tracked frame-by-frame (***C***). ***D***: The speed of larval crawling was averaged across 100 larvae from each genotype (10 videos, each with 10 larvae).

## Discussion

These findings provide novel insights into the role of *sls* in muscle and neuromuscular junction structure and function. We found that *sls* critically mediates proper muscle development, highlighted by our immunohistochemical analyses. Furthermore, muscle impairments in *sls* expression severely disorganizes the NMJ, leading to reduced neuromuscular transduction. Our initial examination of *sls* disruption using ubiquitously expressing muscle drivers validates the critical roles this gene plays in muscle development, highlighted by embryonic lethality resulting from a reduction in *sls* expression. This observation was also recently noted in a *Drosophila* investigation of *sls* [17]. Comparing across the 8 genotypes investigated here, a similar number of eggs were produced from the egg-laying assay. Given that strong, rhythmic muscle contractions are necessary for hatching, it is likely that this prevented the transition from egg to larvae in our assays resulting from severe defects in the development of the NMJ and muscle fibers [28–30]. During vertebrate myogenesis, titin is among the earliest proteins expressed, visible before actin or myosin, and in vertebrate cell culture, developmental perturbations in titin expression profoundly alters muscle structure [12,31]. In *Drosophila, sls* has shown to begin expressing at stage 11 (7 hours after egg laying), and precedes expression of myofilaments, similar to vertebrates [21]. Our screen for Gal4 drivers was a critical step in circumventing the impacts of *sls*-RNAi on embryonic viability, and subsequently enabled a thorough examination of the role of *sls* in *Drosophila*. Using the MF12-Gal4 driver, not only were eggs able to hatch, but animals progressed from egg to adult without any significant impacts on larval, pupal, or adult viability. An examination of whole larvae revealed that disruptions in *sls* expression using MF12-Gal4 did not cause any significant morphological deficits in third-instar larvae. Collectively, *sls* is critically important for the transition from egg to larvae, but the specific cause remains unknown. By limiting the expression of *sls*-RNAi to a single muscle fiber within each abdominal hemisegment, we can circumvent lethality, without affecting the developmental progression from egg to adult. Consequently, we have established the tools necessary for the developmental, molecular, and cellular examination of *sls* in an *in vivo* model.

Upon immunohistochemical examination of individual body-wall muscles from dissected third-instar larvae, the effects of *sls* knockdown on the size and morphology of MF12 were visually striking. Most immediately noticeable were the dramatic 67% decrease in muscle width and 70% reduction in muscle area compared to controls. Since sls-RNAi was selectively expressed in MF12, no other muscle fibers examined were significantly impacted morphologically. Upon closer examination it is improbable the defects in muscle morphology are due to aberrant muscle attachment, as they appear normal for MF12 at each of the abdominal segmental attachment points. Previously in *Drosophila*, an ethylmethane sulfonate (EMS) screen for NMJ disruptions identified novel mutations in *sls* resulting in embryonic lethality [21]. These mutations caused mislocalization, reduction, or complete absence of *sls* expression in embryonic muscles. Most notably these *sls* mutations resulted in a loss of the characteristic striations as well as gross changes in embryonic muscle morphology. In vertebrate skeletal muscle, removal of titin causes muscle atrophy and sarcomeric disassembly [13,32]. In the present study, it is noteworthy that considerable variability in muscle morphology was exhibited both within individual MF12s and between individuals from the MF12>sls genotype. Within individual fibers, most demonstrated a small, and thin phenotype, with areas completely lacking muscle striations, or displaying a spindly appearance. However, there were small, isolated areas along the muscle, particularly the distal ends, where muscles appeared sarcomerically organized. Collectively, *sls* disruption significantly altered gross morphological muscle structure, likely occurring during myogenesis.

During embryogenesis, the somatic muscle system gives rise to the larval musculature [29]. Numerous molecular and cellular processes transform mononucleated muscle precursor cells into multinucleated functional muscles during myogenesis [28]. Individual myoblasts fuse together iteratively to form syncytial myofibers, which requires cytoskeletal elements redistributing and reorganizing many components of the cytoplasmic contents, including myonuclei to regulate internuclear distance [33]. Muscle-cuticle attachments are critical for muscle contraction and ultimately behavior, result from signaling between extending myofibers and apodemes [34]. It is during this time that synaptogenesis and sarcomerogensis are ongoing to establish the contractile machinery underlying excitation-contraction coupling and movement [35]. Two distinct mononucleated myoblasts required for embryonic muscle development are founder cells and fusion competent myoblast, which undergo fusion together [36]. In *Drosophila,* founder cells expressing the immunoglobulin transmembrane protein Dumbfounded recruits the multi-domain adaptor protein Rolling Pebbles which in turn recruits *sls* [21,37,38]. Previous work suggests *sls* is required for myoblast fusion via rearrangement of actin-based cytoskeletal elements in the founder cells. The significant reduction in nuclei observed here supports this idea. *Drosophila sls* is also known to interact with the Z-disc protein mlp84B during development, mediating sarcomeric organization and cellular reorganization during myogenesis [36]. Alterations in this interaction could explain the lack of sarcomeric organization seen in *sls* disrupted animals. The increase in DAPI fluorescence herein is suggestive of increased gene expression, reflective of a compensatory mechanism. Recent work in mammalian skeletal muscle cells sequentially reduced myonuclear numbers and uncovered that myonuclei possess a reserve capacity to support larger functional cytoplasmic volumes in myofibers [39]. They proposed a mechanism of negative correlation between myonuclei number and transcription, which is also supported by our data here [39]. The changes in muscle morphology reported here suggest *sls* disruption impairs myogenesis, likely attributed to impairments in myoblast fusion and cytoplasmic reorganization of cellular elements like nuclei.

Synaptogenesis, where synaptic partner cells recognize one another using a multitude of signaling cues, occurs simultaneously with myogenesis [40]. In *Drosophila,* prior to neuromuscular synapse formation, the embryonic muscles extend dynamic actin-based myopodia, while presynaptic motor neurons extend dynamic growth cones at axon tips searching for extracellular guidance cues [40,41]. Myopodia initially extend randomly, but progressively localize to sites of contact with filopodia from innervating growth cones, in a process of myopodia clustering [41]. Up to 10 distinct motor axons can innervate a single muscle fiber prior to activity-dependent competition leading to single axon retention [42]. Consequently, embryonic muscles are systematically orchestrating chronically occurring dynamic cytoskeletal reorganization throughout myogenesis and synaptogenesis. Here we demonstrate a dramatic consequence of *sls* disruption on the formation and function of neuromuscular synapses. In MF12 from *sls* disrupted animals, NMJ formation largely occurred in the center of muscles, with severely reduced innervation length and bouton number along the muscle. Synapses tended to cluster together, making it difficult to distinguish between the different MN subtypes without tracing back to their separate axonal branches. While significant impacts were observed for both MN subtypes, similar effects indicate MN subtype-specific synaptic targeting precedes synaptogenesis and indicates an earlier developmentally aberrant process.

Within individual boutons, neurotransmitters (NTs) contained within synaptic vesicles (SVs) fuse at AZs, at the core of which is the electron dense scaffolding protein, brp, responsible for SV clustering and organization of the AZ components and fusion machinery (e.g. calcium channels, [43,44]. AZ assembly has been heavily investigated in *Drosophila* [45,46]. AZ brp levels strongly correlate with SV release probability, and their density varies considerably in responses to presynaptic structural changes [47–49]. Recently, much attention has been focused on the role AZ proteins in homeostatic plasticity in compensating for altered SV release [50–52]). The significant reduction in synaptic efficacy resulting from *sls* knockdown here may also trigger these pathways, resulting in changes in AZ structure. Thus, the changes in AZ brp density we observed could reflect developmental structural changes resulting from aberrant synaptogenesis, or a more dynamic processes ongoing mediating mechanisms of SV release. Postsynaptic ionotropic glutamate receptors depolarize muscle cells in response to SV fusion and NT release. These heteromeric tetramers are composed of GluRIII, GluRIID, GluRIIE, and a variable fourth subunit, either GluRIIA, or GluRIIB [53]. Our GluRIII immunostains did not reveal any differences in postsynaptic density or AZ density. This provides evidence that postsynaptic receptor complex formation is occurring normally, likely in response to presynaptic AZ seeding [53]. However, GluRIIA/B are known to be dynamically regulated during synapse maturation, and the two exhibit different ionic conductances [47,54]. Postsynaptic GluRIII density has been shown to remain unchanged, even when presynaptic AZ proteins were altered significantly [55,56]. To our knowledge, this is the first time *sls*, a putative homologous structure of human titin, has been demonstrated to significantly impact synapse formation and synaptic communication.

As muscle contractility is based on a complex protein structure of repeating individual contractile units of nearly crystalline order, an examination of the ultrastructural changes in MF12 following *sls* disruption was logical. One of the long-seeded postulates for the structural role of *sls*/titin is to maintain A-band stability during and after muscle contraction [13,57]. Within the actin and myosin (two-filament) cross-bridge model of contraction, sarcomeres and half-sarcomeres are unstable [58,59]. Small differences caused by stochastic interactions of cross-bridges with actin will lead to an initially centered myosin filament to be displaced leading to unstable and uneven overlap between actin and myosin within each half-sarcomere. This structural asymmetry and uneven interaction between the two filaments will lead to A-band drift, and unequal force generation on either side of the M-line [13]. Indeed, this A-band shift has been observed following prolonged activation of rabbit psoas muscles when titin’s contribution was effectively eliminated [60]. In our experimental setup, wandering third-instar larvae were isolated, dissected and fixed at varying phases of peristaltic locomotion, i.e. forward vs reverse, contraction vs relaxation phases of crawling. Given the highly variable nature of larval crawling, it is logical to predict that if *sls* was responsible for A-band stability, then the results of *sls* disruption in our model would be highly variable I-band lengths, which is indeed what we observed. Previously it has been suggested that the length of the so-called super-repeats domains within mouse titin, matches the 43-nm distance of the myosin heads in the C-zone, a region of each half-sarcomere flanking the A-band containing the myosin binding sites [61]. Recent work in the mouse demonstrated that deletion of a subset of these super-repeats resulted in shorter A-bands [62]. Our results revealed a doubling of the average A-band length following *sls* knockdown; therefore, it seems plausible that *sls* is working as a molecular ruler to regulate A-band length. It is noteworthy that in *Drosophila,* Loreau et al, (2023), revealed that Projectin decorates the entire thick filament of larval body-wall muscle sarcomeres, while *sls* only interacts with the lateral-most aspect of myosin. Therefore, Projectin is uniquely positioned to interact with *sls* in its role of molecular ruler for thick-filament length.

Another putative role for *sls*/titin is as a stretch sensor which activates super-relaxed cross-bridges, and therefore provides a mechanism for compensating for reduced actin-myosin overlap (elongated sarcomere) by increasing the density of cross-bridges at locations of myofilament overlap [63]. *Drosophila* body-wall muscles are supercontractile, able to contract well below 50% of resting length [64]. *Drosophila sls* could serve a role in regulating cross-bridge kinetics in sarcomeres of different lengths to regulate stability [63]. Furthermore, since adjacent half-sarcomeres overlap in the Z-disc and M-band regions, there is the capacity for continuous structural and functional (force) transmission, providing a mechanism for *sls*-mediated orchestration across sarcomeres [65]. This would facilitate synchronicity across the entire muscle, making *sls* an integral aspect of the contraction generating machinery by transmitting myofibril-based forces along adjacent sarcomeres.

The role of titin in active, passive, and residual force enhancement is by far the most well characterized and discussed (reviewed: [8–10]). We looked beyond the effects of *sls* on muscle contractile properties and examined changes in muscle contractile behavior, via a crawling assay. Surprisingly, reducing the expression of *sls* in a single muscle fiber within each abdominal hemisegment was sufficient to significantly reduce larval crawling behavior. Our previous work demonstrated that MFs 6, 7, 12, and 13 contribute most substantially to longitudinal force production underlying motivated forward peristalsis along the ventral substrate [66]. Previously, we serially ablated these fibers and found that muscles 12/13 contributed ∼30% of the total longitudinal force production, and likely explains the significantly reduced larval crawling velocity shown here [26,66]. In addition to these putative roles of titin/*sls*, this elastic-protein has also been suggested to serve a multitude of other roles via its interactions with sarcomeric and non-sarcomeric proteins. These functions are incredibly diverse and include the Blaschko effect (a.k.a catch-tension), mechanosensory, signaling hub, target of proteostasis mechanisms, substrate for calcium binding, and posttranslational modifications (phosphorylation or oxidation) regulating stiffness [8–10,60]. Given the robust toolkit assembled herein along with those within the *Drosophila* community, many of these processes can, and should be investigated in future studies. Here we have demonstrated profound effects for *sls* on muscle structure, function, neuromuscular transduction, and ultimately, locomotory behavior.

## Materials and Methods

### Husbandry

*Drosophila melanogaster* were cultured on standard medium at 25°C, at constant humidity, and in a 12:12 light: dark cycle. Genotypes used in this study include the following: UAS-sls1-RNAi ([31538, Bloomington Drosophila Resource Center (BDRC)]; UAS-sls2-RNAi (31539, BDRC); GAL4-Muscle Fiber 12 (91395, BDRC); GAL4-Mef2 DICER(X) (25756, BDRC); UAS-GFP (1522, BDRC); MHC-Gal4 (55133, BDRC).

### Immunohistochemistry

Wandering third-instar larvae were dissected in hemolymph-like saline HL3.1 solution with the following composition (in mM): 70 NaCl, 5 KCl, 0 CaCl2, 4 MgCl2, 10 NaHCO3, 5 trehalose, 115 sucrose, and 5 HEPES, pH 7.2 [67]. The larvae were fixed for three minutes in 4% paraformaldehyde and subsequently washed in phosphate-buffered saline (PBS), with 0.05% Triton X-100 (PBST). Animals were then incubated with primary antibodies in PBST at room temperature for 2 hours and washed three times for 10 min in PBST. Secondary antibodies were added to fresh PBST solution and were incubated at 4°C overnight. Finally, larvae were washed three times for 10 minutes in PBS and then mounted in medium containing DAPI (ab104139). Antibodies used for this study include the following: mouse anti-Myosin, 1:500 [stock #EB165, Developmental Studies Hybridoma Bank (DSHB)]; rat anti-Sallamus (*sls*), 1:500 (stock #4F3, DSHB); mouse anti-brp (Bruchpilot), 1:500; rabbit GluR3 1:2000 (Gift, Troy Littleton); goat anti-mouse Alexa Fluor 578, 1:500 (catalog #A-16071; Thermo Fisher Scientific); and phalloidin-conjugated Alexa Fluor 488, 1:4000 (Thermo Fisher Scientific). Immunoreactive proteins were imaged on a Nikon fluorescence microscope (Nikon Instruments Inc) using 20 or 60x magnification and processed using Nikon Elements software.

### Morphological Measurements

Muscle length and width measurements were taken using an upright standard dissecting microscope with an objective containing a reticle from 20 randomly selected wandering third-instar larvae for each control and mutant line. The larvae were rinsed in DI water prior to being measured. Larvae were pinned and stretched until abdominal contractions were no longer observed. Area was calculated using as the product of length times width. *Muscle specific analyses:* Muscle length was measured using the distance measurement feature on the Nikon Elements software. Length was measured at the midpoint of each endline of the muscle. Width was gathered using the same Nikon tool from the middle of the muscle fiber. Muscle area was calculated as the product of both. Muscle sarcomere length was measured using the intensity profile function in Nikon elements software to produce a sinusoidal graph of anti-actin fluorescence intensity. Then, sarcomere length was measured as one sinusoidal wavelength. I-band was also collected using the anti-actin stain and was measured as the width of each peak measured from the first substantial dip in fluorescence. A-band was measured using an anti-myosin stain, using the width of each peak. The *sls*-band was measured similarly to the I- and A-band measurement but using an anti-*sls* stain. Fluorescent intensity was measured by placing two identical boxes on muscle 12 and 13 of all three genotypes. Fluorescence intensity (AU) was exported using Nikon elements software. DAPI mounting media (ab104139) was used when mounting fixed dissections on microscope slides. Nuclei area was measured from overlayed images of DAPI and anti-actin using a region of interest (ROI) measuring tool on Nikon elements. Subsequent fluorescence intensity, number, and area of each ROI was then exported to Microsoft Excel. Neuromuscular junction analysis was performed using anti-brp or anti-GluR3 primary antibodies, and fluorescent secondaries. ROIs were then placed using the Nikon elements tool in a similar manner to the DAPI analysis. Bouton average active zone fluorescence, bouton number, and bouton area were subsequently exported to excel. Bouton density was calculated through taking the area of each bouton and manually counting the number of active zone puncta. Density was calculated as bouton area/puncta number. ***Fecundity.*** For each genotype assayed: three males and three females were isolated upon eclosion and remained in a vial until reaching day three when they were transferred onto Petri dishes containing grape agar and yeast paste. Flies were left for 24 hours on grape agar dishes, and the number of eggs were recorded and organized in rows of ten. Twenty-four hours later, the number of hatched eggs were recorded, and larvae were transferred to clean food vials to be later checked for pupation. The grape agar was made with 25% grape juice, 75% water, 3% agar, and 0.3% sucrose. The yeast paste was a mixture of DI water and active dry yeast with a consistency of creamy peanut butter.

### Electrophysiology

Wandering third-instar larvae dissected in HL3.1 (0.3mM Ca^2+^) were dissected and pinned dorsal side up, all nerves emerging from the ventral nerve cord were severed, and the brain and ventral nerve cord were removed [67]. Severed nerve branches were electrically stimulated with a suction electrode connected to a Master 8 stimulator (A.M.P.I.). Intracellular voltage recordings were obtained using an Axoclamp 2B amplifier (molecular devices) and digitized using a minidigi 1b (molecular devices). Signals were acquired using Clampex and analyzed using Clampfit, MiniAnalysis, Microsoft Excel, and GraphPad Prism.

### Larval Crawling

Wandering third-instar larvae were removed from the sides of culture vials, washed seven times in deionized water and placed on Petri dishes containing 1% agar. For each recording 10 larvae were placed in the center of a dish. The dishes containing larvae to be recorded were placed within a custom-made black opaque box, creating a completely dark environment. The larvae were recorded using an infrared camera (FLiR blackfly), which connected to a standard computer. Each video was recorded using SpinView (FLIR) software for a length of five minutes. The videos were converted to interoperable master format and subsequently analyzed by CTRAX. Files were exported from CTRAX as a CSV file readable by excel. CTRAX extracted x and y coordinates of each larva and the angle of ellipse, or heading, in radians. Using excel, each x and y coordinate of each larva was translated into pixels, and values such as displacement, velocity over a 0.5 second interval, total distance travelled, and angular velocity were found.

### Statistical analysis

Prism software (version 10.1.0; GraphPad Software) was used for statistical analysis. Appropriate statistical metrics were performed for each dataset which was included in the results section along with the F, and P statistics. Statistical comparisons were made with controls unless noted. Appropriate sample size was determined using a normality test. Data are presented as the mean ± SEM (*p< 0.05, **p< 0.01, ***p< 0.001, n.s. = not significant).

## Notes

### Competing Interest Statement

The authors have declared no competing interest.

